# Competitive Inhibition of the Nipah Virus Matrix Protein by Conivaptan: From Binding Pocket Dynamics to Protein-Membrane Interactions

**DOI:** 10.64898/2026.08.02.742314

**Authors:** Amar Prasad Kar, Atanu Maity, Ranjit Prasad Bahadur

**Affiliations:** Bioinformatics Center, Department of Bioscience and Biotechnology, Indian Institute of Technology Kharagpur, Kharagpur-721302, India; Computational Structural Biology Laboratory, Department of Bioscience and Biotechnology, Indian Institute of Technology Kharagpur, Kharagpur-721302, India

**Keywords:** Nipah Virus, Matrix protein, Ensemble docking, Drug repurposing, Protein-membrane interaction

## Abstract

The recurrent outbreaks of Nipah virus since its first emergence have intensified efforts to identify effective antiviral therapeutics. The matrix protein is a key structural protein that maintains virion architecture and plays a crucial role in anchoring the virus to the host plasma membrane. Binding of phosphatidylinositol-4,5-bisphosphate (PIP2) to the matrix protein promotes conformational changes that facilitate membrane association and initiate viral assembly. Therefore, an effective small-molecule inhibitor should not only compete with PIP2 for binding to the Nipah virus matrix protein (NiVM) but also disrupt its interaction with the host membrane. Using virtual screening of FDA-approved drugs combined with ensemble docking, we identified Conivaptan as a potential inhibitor of the Nipah virus matrix protein. The multivalent interactions of Conivaptan establish stable binding within the PIP2-binding pocket while simultaneously rewiring protein-membrane interactions. In the presence of PIP2, NiVM anchors to the membrane through multiple charged residues, resulting in pronounced local bilayer deformation, an early signature of membrane remodeling. In contrast, Conivaptan binding weakens membrane anchoring and preserves a more ordered and compact bilayer, thereby suppressing membrane deformation. Together, these findings identify Conivaptan as a promising repurposable inhibitor of the Nipah virus matrix protein and provide a mechanistic framework for targeting virus-membrane interactions during viral assembly.

## 1. Introduction

Nipah virus (NiV), a member of the *Henipavirus* genus within the *Paramyxoviridae* family^1,2^, is an emerging zoonotic pathogen that has caused recurrent outbreaks in South and Southeast Asia in recent years^3^. In humans, infection causes severe respiratory illness and can often progress to deadly encephalitis, with death rates ranging from 40% to over 70% across outbreaks^4,5^. Due to its high pathogenicity, broad host range, and capacity for human-to-human transmission, NiV is designated a Biosafety Level-4 (BSL-4) agent and listed by the World Health Organization (WHO) as a priority pathogen for accelerated research into countermeasures^6–8^. Despite its significant public health impact, no licensed vaccines or therapeutics are currently available, underscoring the need for the development of effective interventions^5,9,10^.

The Nipah virus genome encodes six structural proteins which are Glycoprotein, Matrix protein, Fusion protein, Nucleoprotein, Phosphoprotein, and RNA-directed RNA polymerase.^11^ Among them, the matrix protein plays the central role in viral assembly and budding^12^. Recent studies have revealed that Matrix protein-membrane interaction is sensitive to the presence of phosphatidylinositol-4,5-bisphosphate (PIP2). The presence of PIP2 along with phosphatidylserine (PS) facilitates Matrix protein recruitment to the plasma membrane, promoting lattice formation. This induces membrane curvature necessary for viral budding^13^. This PIP2-dependent control of Matrix protein localization and assembly underscores its importance in NiV infectivity and reinforces its potential as an antiviral target. Nipah virus Matrix protein interacts with plasma membrane in a dimeric state. The structural characterization of Matrix protein in the presence (PDB ID: 7SKU) and absence of PIP2 (PDB ID: 7SKT) reveals a large conformational change of its C-terminal residues^13^. The last 24 residues of each monomer of the apo dimer are closely packed into the structural core by forming intra-monomer interaction (Figure 1). Whereas, in the PIP2-bound dimer, the tail is extended away from the core and interacts with the other monomer resulting in a domain-swapped dimer. The PIP2 binding pocket is occupied by a small helical segment termed as alpha-2 helix in the apo dimer. Norris *et al.* have shown that PIP2 binding reshapes the electrostatic surface of the Matrix protein, triggering conformational changes.^13^ These conformational changes facilitate matrix protein dimerization and membrane deformation, both of which are essential for efficient viral budding. Since PIP2 binding governs the structural state of the matrix dimer, the corresponding interaction surface represents a rational target for competitive inhibition. PIP2 engages the matrix protein through localized electrostatic interactions, indicating that activation depends on specific headgroup binding rather than diffuse membrane contact. As this interaction relies on localized charge complementarity, it may be competitively disrupted by small molecules engaging the same residues. This provides a structural rationale for targeting the PIP2-interacting region in a competitive inhibitor design strategy.^14^

**Figure 1:**
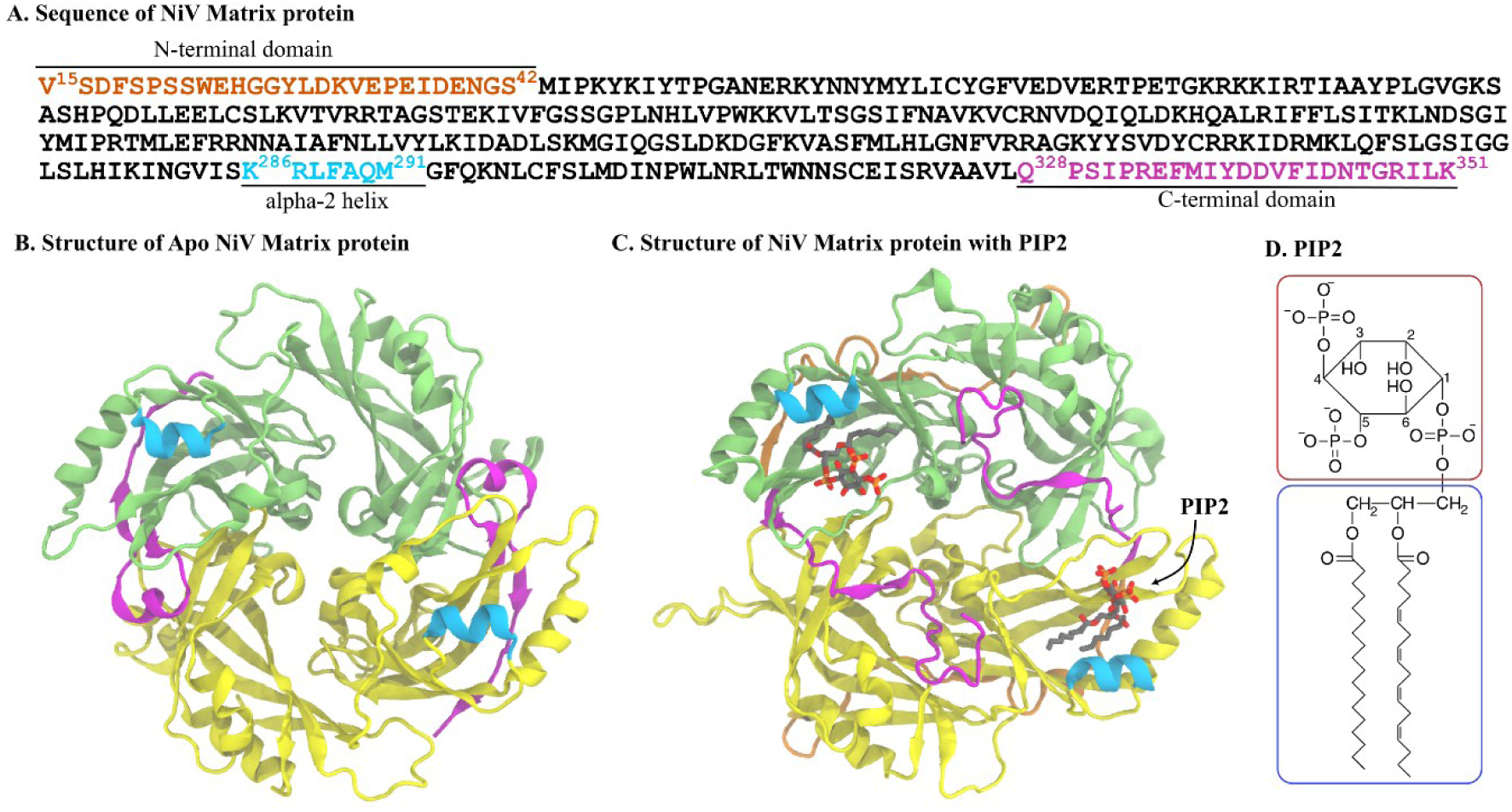
Structures of Nipah virus matrix protein (NiVM). A) Sequence of the monomeric unit of NiVM. 3D crystallographic structure of NiVM in B) apo (PDB ID: 7SKT) and PIP2-bound (PDB ID: 7SKU) states. Two monomers are colored in green and yellow. N-terminal end, C-terminal end and alpha2 helix are shown in orange, magenta and cyan, respectively. PIP2 is represented as stick. D) Molecular structure of PIP2. Polar head group and non-polar acyl chains of the marked with red and blue rectangle respectively.

Despite these advances, important aspects of PIP2-mediated regulation of the Nipah virus matrix protein remain incompletely understood.^13,15^ In particular, static structural models provide limited insight into the dynamic nature of PIP2 recognition and the conformational flexibility of the binding pocket, both of which may influence the identification of competitive inhibitors. Furthermore, the effect of small-molecule binding on the membrane-associated function of the matrix protein has not been systematically evaluated. Addressing these limitations requires approaches that integrate protein conformational dynamics with membrane-associated functional analyses to enable a more comprehensive assessment of candidate inhibitors.

In the present study, we combined ensemble-based virtual screening and docking, and membrane simulations to identify FDA-approved compounds capable of competitively targeting the PIP2-binding pocket of the Nipah virus matrix protein. Our analyses identify Conivaptan as a promising repurposable inhibitor with a dual mechanism of inhibition. Conivaptan not only exhibits stable binding within the PIP2-binding pocket but also perturbs the protein-membrane interactions of NiVM that are required for membrane remodeling. By integrating conformational heterogeneity of binding pocket with the dynamics of protein-membrane interactions, this study provides mechanistic insights into matrix protein inhibition and establishes a framework for the rational design and evaluation of antiviral therapeutics targeting Nipah virus.

## 2. Material and methods

### 2.1. Modelling of apo and PIP2-bound matrix protein

The Apo form (PDB ID: 7SKT) of matrix protein and its complex with PIP2 (PDB ID: 7SKU) were obtained from the RCSB PDB^13^. Both structures are crystalized as dimer which is also their biologically relevant form. The missing residues were modelled using SWISS-MODEL^16^. Crystal water and other non-essential heteroatoms (except PIP2) were removed from the structures using PyMOL^17^. Another system was prepared by removing the PIP2 molecule from the complex structure to check the binding pocket flexibility.

### 2.2. Molecular dynamics simulations of Apo and PIP2-bound NiV-Matrix Protein

Each system was solvated in a waterbox containing an adequate number of TIP3P water molecules^18^. The dimensions of the boxes were determined by maintaining a minimum of 1 nm distance between the protein surface and the edge of the waterbox. Appropriate numbers of sodium and chloride ions were added to achieve 0.15M salt concentration after neutralizing the system to mimic the physiological condition. CHARMM36 all-atom force field for protein^19^ along with CGENFF^20^ force field for small molecules were used to model the systems. Each system was energy minimized for 2000 steps using ABNR followed by a 1000 step SD energy minimization algorithm to remove unfavourable contacts and steric clashes. Systems were equilibrated in NVT ensemble for 1 ns followed by 5 ns equilibration in NPT ensemble. V-rescale thermostat^21^ was used for temperature coupling to maintain 300K temperature during NVT and NPT equilibration. Parrinello-Rahman barostat^22^ was used for pressure coupling to maintain a pressure of 1 atmosphere during NPT equilibration. Equilibration dynamics was followed by 1 µs production for each system. All simulations were run using GROMACS 2024.2^23^. One fs timestep was used for NVT equilibration. LINCS^24^ algorithm was used to constrain all bonds with hydrogen atoms during NPT equilibration and production dynamics to allow 2 fs timestep. A 1.0 nm cut-off distance was used for the calculation of short-range electrostatics and van der Waals interactions. Long-range electrostatic interactions were taken care using the Particle-Mesh Ewald (PME)^25^ summation method with periodic boundary condition. Coordinates and velocities were saved at an interval of 10 fs during the production dynamics. Each simulation was repeated thrice starting from the NVT equilibration to ensure sufficient sampling.

### 2.3. Preparation of target protein and FDA-approved ligands

To account for the flexibility of the PIP2 binding pocket, an ensemble docking approach was adopted to target the pocket with small molecule inhibitor. The trajectory of PIP2-bound NiV MP was clustered using *gmx_cluster* based on the root mean square deviation (RMSD) of the PIP2 binding pocket amino acids. A RMSD cutoff of 1.5 Ả was used after removing PIP2 from the binding pocket. Representative structures from the three most populated clusters were selected for docking. AutoDockTools 1.5.7^26^ was used to add polar hydrogens and assign Gasteiger charges to each structure. Docking was carried out with AutoDock Vina 1.2.5^27^. For all three receptors, a 30×30×30 Å³ grid box was centered on the PIP2-binding pocket. The grid boxes were centered at cartesian coordinates (-0.891, -0.832, -0.956), (-1.395, -0.636, -1.731) and (-0.891, -0.832, -2.466) for the three receptors. Same docking parameters (exhaustiveness = 20, num_modes = 20, energy_range = 1 kcal/mol) were used for the three receptors. A total of 1205 FDA approved drug molecules were selected as ligands for virtual screening and their structures were obtained from Pubchem^28^. The screening of ligands including ligand preparation, docking, scoring and complex structure generation was handled using a perl-based pipeline and in-house python scripts. The top 10 ranked binders from the three receptors were compared and the common ligand that appeared within the top ten binders across the three receptor conformations was chosen for further analyses. The complex between the chosen ligand with the receptor from cluster1 was selected as the inhibitor bound NiVM. This structure was further refined and simulated following section 2.2.

### 2.4. Simulation of membrane-embedded matrix proteins

To investigate the membrane interaction of the Nipah virus matrix (NiV-M) protein in its major functional states, we simulated NiVM in apo and small molecule-bound form embedded in a plasma membrane bilayer. The composition of the two leaflets of the bilayer was chosen to model the asymmetric plasma membrane^29^. The outer leaflet contains 205 lipid molecules including 88 POPC, 15 POPE, 6 POPS, 6 PIP2, 47 PSM and 43 cholesterols whereas the inner leaflet has 208 lipid molecules (21 POPC, 75 POPE, 32 POPS, 28 PIP2, 9 PSM and 43 cholesterols). CHARMM-GUI membrane builder^30^ was used to build the bilayer assemblies and to embed the protein on the bilayer. The protein was placed closed to the inner leaflet with the PIP2 pocket facing the lipid headgroup. TIP3P water molecules were added to both sides of the bilayer to hydrate the bilayer. The thickness of water layer at the outer leaflet was 3 nm and it was 7 nm on the inner side of the bilayer containing the protein (Figure S1). An adequate number of counter ions were added to achieve 0.15M ion concentration after neutralizing the total charge of the systems. The assembly was minimized in three steps using positional restraints to specific atoms. In the first step the waters, ions and the heavy atoms of the protein molecule and phosphate atoms of the membrane were restrained to minimize the remaining system for 5000 steps. In the second step, restraints are removed from water and ions keeping rest of the component same for another 5000 steps of energy minimization. In the final step, all restraints were removed and the system was minimized for additional 5000 steps. The systems were then equilibrated through a six-step protocol with gradually decreasing positional and dihedral restraints. The first three equilibration stages were performed for 10 ns each, followed by three additional stages of 20 ns each. Initially, strong harmonic restraints were applied to the protein backbone (4000 kJ mol⁻¹ nm⁻²), protein side chains (2000 kJ mol⁻¹ nm⁻²), lipid headgroups (1000 kJ mol⁻¹ nm⁻²), and protein dihedral angles (1000 kJ mol⁻¹). These restraints were gradually reduced over six equilibration stages. These equilibration steps were followed by a 20ns NPT equilibration phase to ensure complete relaxation of the membrane environment before production simulations. One microsecond production dynamics was performed for all three systems: Bilayer_NiVM, Bilayer_NiVM_PIP2 and Bilayer_NiVM_Conivaptan. The simulations were triplicated starting from the NVT equilibration. An additional 1 microsecond simulation of pure lipid bilayer was performed as a control system. Simulations were performed in GROMACS using appropriate CHARMM forcefield for protein and lipids and small molecules. The use of thermostat and barostat and the target temperature and pressure were same as mentioned in section 2.2. Other simulation parameters and conditions were similar to the methodology section 2.2. All simulated systems are listed in Table 1.

**Table 1:** Details of the systems simulated.

| System identifier | Protein | Membrane | Small molecule | Simulation length |
| --- | --- | --- | --- | --- |
| NiVM | Nipah virus Matrix protein (modelled based on 7SKT) | — | — | 1 μs |
| pseudo unbound | Nipah virus Matrix protein (modelled based on 7SKU) | — | — | 1 μs |
| NiVM_PIP2 | NiVM | — | PIP2 | 1 μs |
| NiVM_Conivaptan | NiVM | — | Conivaptan | 500 ns |
| Bilayer_NiVM | NiVM | asymmetric bilayer | — | 1 μs |
| Bilayer_NiVM_PIP2 | NiVM | asymmetric bilayer | PIP2 | 3 x 1 μs |
| Bilayer_NiVM_Conivaptan | NiVM | asymmetric bilayer | Conivaptan | 3 x 1 μs |
| Bilayer | — | asymmetric bilayer | — | 1 μs |
| Total | | | | 11.5 μs |

### 2.5 Calculation of membrane physical properties

The basic structural parameters were calculated using in-built utilities of GROMACS. Membrane thickness was calculated using a Perl-based interface Gridmat-MD^31^. Membrane thickness was estimated at each x-y grid as the average distance between a selected head-group atom of lipid molecules from the upper and lower leaflet. Phosphorus atom of the phosphate head group was considered for all lipid types except cholesterol The values were collected considering grids of length 1 Å in both x and y-direction and averaged over the simulation time. Acyl chain order parameter was calculated using an in-built module of GROMACS. Order parameter was calculated separately for palmitoyl and oleoyl chains and averaged over all lipid types at different carbon position using the following formula -

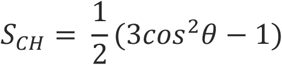

where *θ* is the angle of the resultant of two C-H bonds with respect to the membrane normal. MDTraj^32^ was used for calculating hydrogen bonds and contacts between the protein-protein and protein-lipid.

## 3. Results

### 3.1 Conformational rearrangement of NiVM upon PIP2 binding

The binding pocket residues of NiVM comprised of both hydrophobic (L239, M262, L264, I278, L348) and hydrophilic residues (Q195, N241, R198, Q291). On the other hand, PIP2 has two highly polar phosphate groups and two non-polar acyl chains. This facilitates the possibility of both polar and non-polar protein-ligand interactions. Polar interaction was monitored as the number of hydrogen bonds between the polar atoms of PIP2 and hydrophilic residues of the binding pocket. The non-polar interaction was quantified as the number of contacts between non-polar heavy atoms of PIP2 and the hydrophobic amino acids of the binding pocket. The polar interaction is governed by the hydrogen bonds formed between the hydroxyl group of PIP2 and Q291 in the binding pocket. This hydrogen bond is persistent over 40% of the simulation time (Figure 2A). The average number of hydrophobic contacts between PIP2 and the binding pocket residues is 17 (Figure 2B). The hydrophobic residues of the binding pocket are shown in yellow (Figure 2A(ii)). The strong hydrophobic and polar interaction helps PIP2 to be deeply inserted into the binding pocket (Figure 2C).

**Figure 2:**
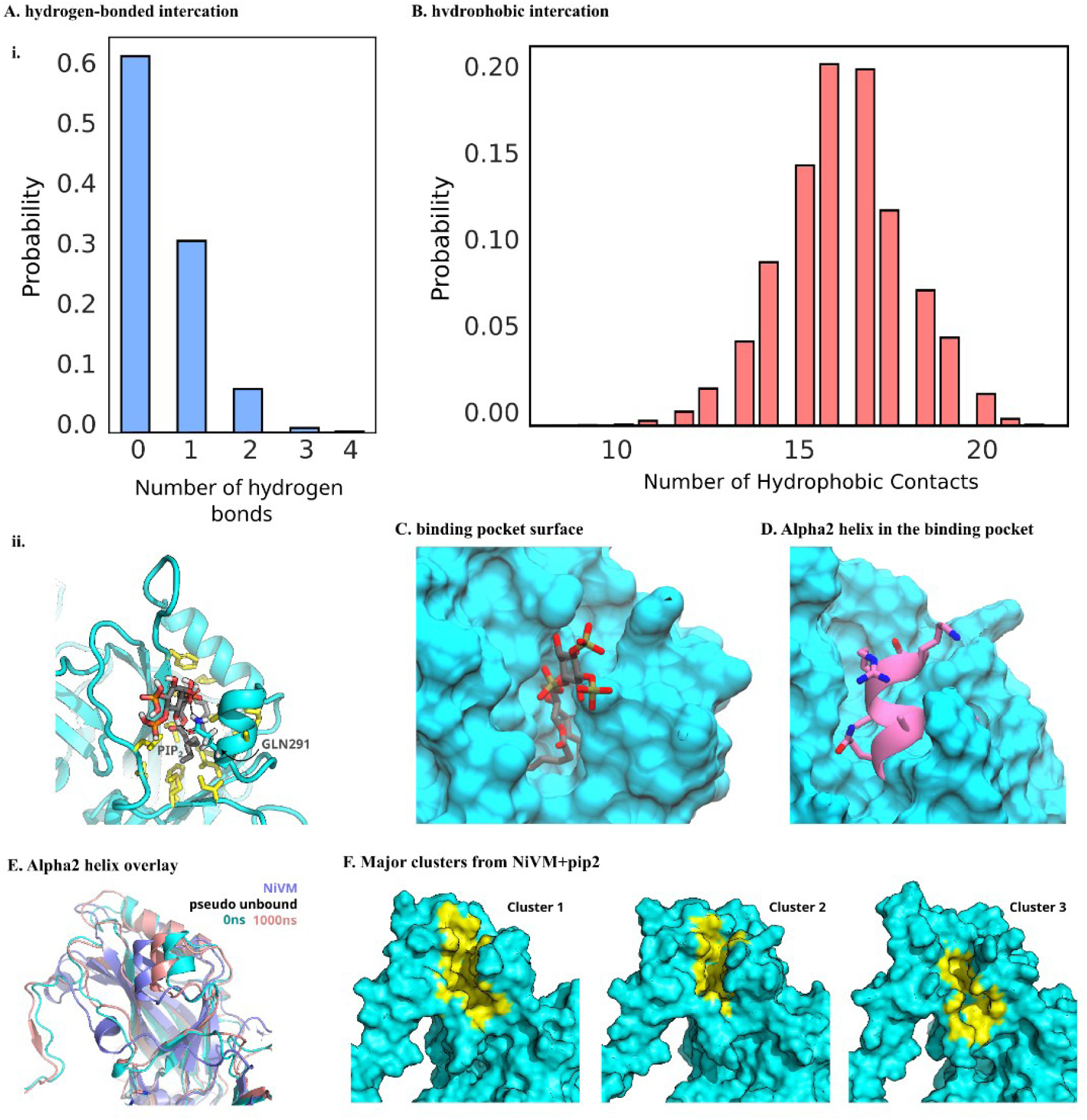
Dynamic nature of PIP2 binding and structural plasticity of the NiVM PIP2-binding pocket. (A) (i) Distribution of hydrogen bonds formed between PIP2 and NiVM (ii) Representative binding mode of PIP2 in the NiVM binding pocket illustrating the interacting residues. (B) Distribution of hydrophobic contacts between PIP2 and NiVM pocket residues during the simulation. (C) Surface representation of the PIP2-binding pocket showing the orientation of PIP2 within the cavity. (D) Conformation of the α2 helix occupying the PIP2-binding pocket. (E) Superposition of the α2 helix from the pseudo-unbound structure, the initial simulation frame (0 ns), and the final simulation frame (1000 ns), illustrating conformational rearrangements during the simulation. (F) Surface representations of the three most populated conformational clusters obtained from clustering analysis of the NiVM– PIP2 trajectory, highlighting the structural heterogeneity of the PIP2-binding pocket

In the absence of PIP_2_, the binding pocket is occupied by the residues of alpha2 helix (Figure 2D). The side chain of Gln291 of alpha2 helix is involved in hydrogen-bonded interactions with the backbone of Asn241 present in the beta sheet of the binding pocket. On the other hand, Met292 and Leu288 form hydrophobic contacts with hydrophobic residues of the binding pocket. The differences in the interaction profile of PIP2-bound and apo NiVM indicates a large-scale conformational rearrangement of the binding pocket and surrounding residues.

To find whether the structural rearrangement observed in the PIP2-bound NiVM are irreversible, we prepared an unbound NiVM dimer by removing PIP2 from the binding pocket and called it ‘pseudo unbound’. During the simulation, there are considerable rearrangement of the binding pocket residues. At the end of 1 µs, the alpha2 helix shows a propensity to adopt apo-like conformations partially covering the binding pocket (Figure 2E). The adaptability of the PIP2 binding pocket residues poses a challenge to the conventional single-target drug-designing approach and highlights the importance of an ensemble-based approach.

### 3.2 Binding-pocket clustering and ensemble docking

The trajectories of PIP2-bound NiVM were clustered using root mean square deviation (RMSD) of binding pocket residues to identify the major clusters. Clustering yields six major clusters which cumulatively cover ∼60% population of the NiVM_PIP2 trajectory (Figure S2). The binding pocket conformations vary significantly in terms of the shape of the binding pocket and the interaction of PIP2 with binding pocket residues. The representative structures of the three major clusters are shown in Figure 2F. These most populated conformations were selected as representative receptor structures for subsequent ensemble docking. The orientation of the ligand in the three major clusters are different to complement the pocket conformation (Figure 3A). This ensures that ligand screening incorporates the conformational variation emerging from PIP2-binding. All of the 1205 FDA-approved small molecules were screened against these three targets and ranked according to docking score. The dock score of the top 10 molecules ranges from -8.7 kcal/mol to -11.23 kcal/mol and in the three docked complexes (Table S1). The top ten molecules docked to the most populated cluster (cluster 1) are listed in Table 2 and shown in Figure S3. The top-ranked molecules included several clinically approved compounds such as Lumacaftor (PubChem CID: 16678941), Lomitapide (PubChem CID: 9853053), Midostaurin (PubChem CID: 9829523), Dutasteride (PubChem CID: 6918296), Netupitant (PubChem CID: 6451149), Lapatinib (PubChem CID: 208908), Telmisartan (PubChem CID: 65999), Lurasidone (PubChem CID: 213046), and Nilotinib (PubChem CID: 644241), exhibiting predicted binding energies in the range between -9.48 kcal/mol and -9.88 kcal/mol.

**Figure 3:**
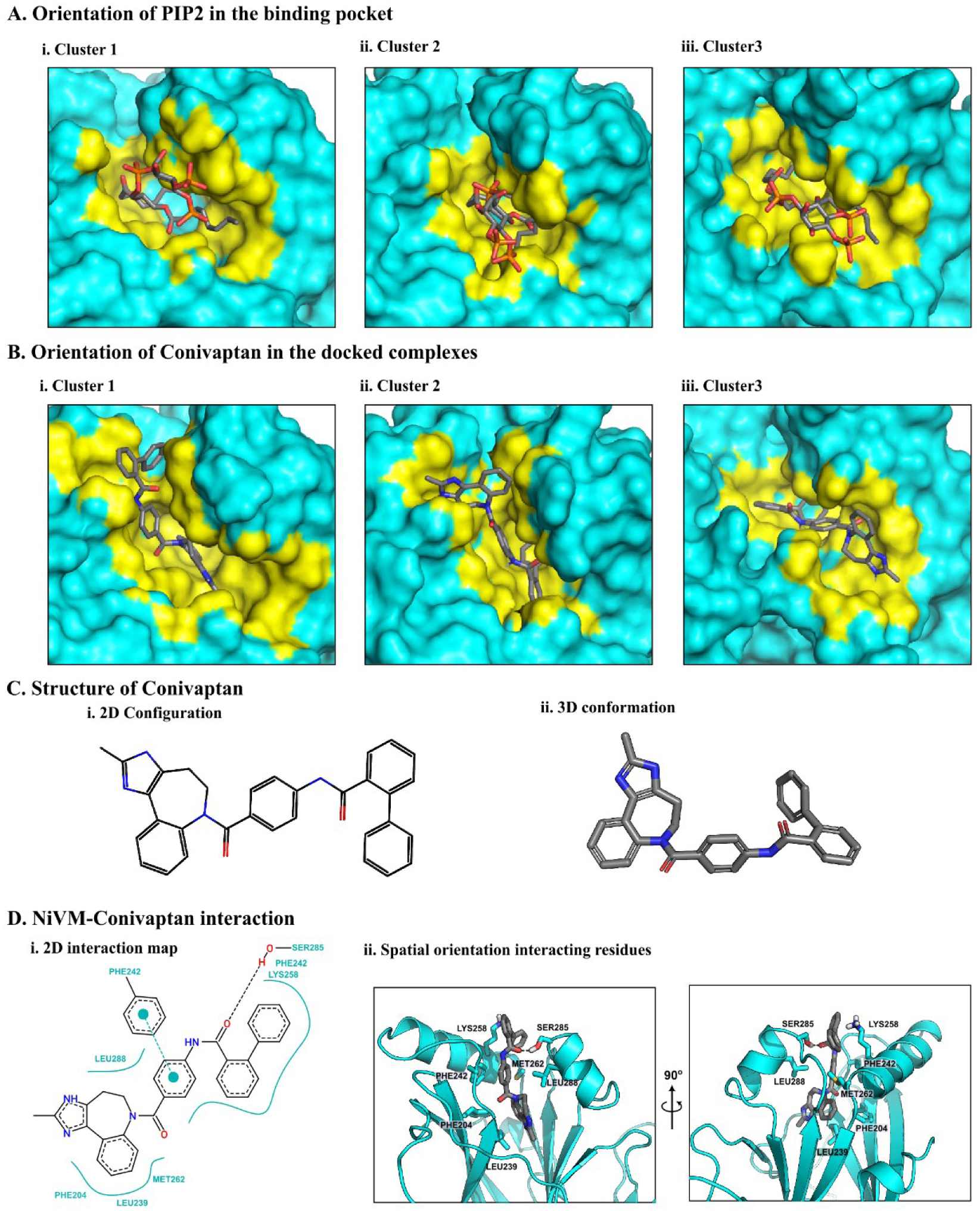
Representative conformations of the NiVM PIP2-binding pocket and docking of Conivaptan to the major conformational clusters obtained from molecular dynamics simulations. (A) Orientation of PIP2 in the three most populated clusters (Cluster 1–3) of the NiVM binding pocket. The protein surface is shown in cyan, while the PIP2-binding pocket is highlighted in yellow. (B) Binding orientations of Conivaptan docked into the representative structures of Clusters 1–3, demonstrating favorable accommodation of the ligand across the different pocket conformations. (C) Chemical structure of Conivaptan showing (i) the two-dimensional chemical structure and (ii) the optimized three-dimensional conformation. (D) Predicted interactions between Conivaptan and NiVM. (i) 2D interaction map showing hydrogen bonds and hydrophobic contacts between Conivaptan and the binding pocket residues. (ii) 3D representation of the docked complex highlighting the spatial arrangement of interacting residues around Conivaptan from two orthogonal orientations.

**Table 2:** Details of top 10 FDA-aproved drugs obtained from virtual screening targeting binding pocket of cluster1.

| Rank | Ligand Name | PubChem CID | Predicted Binding Affinity (kcal/mol) |
| --- | --- | --- | --- |
| 1 | Conivaptan | 151171 | -11.23 |
| 2 | Telmisartan | 65999 | -9.877 |
| 3 | Midostaurin | 9829523 | -9.813 |
| 4 | Dutasteride | 6918296 | -9.782 |
| 5 | Lurasidone | 213046 | -9.733 |
| 6 | Lapatinib | 208908 | -9.602 |
| 7 | Lomitapide | 9853053 | -9.595 |
| 8 | Nilotinib | 644241 | -9.528 |
| 9 | Lumacaftor | 16678941 | -9.553 |
| 10 | Netupitant | 6451149 | -9.487 |

Among the compounds binding to the three clusters, Conivaptan (PubChem CID: 151171) is common (Figure 3B) with high binding affinity. Conivaptan (Figure 3C) is an FDA-approved, non-peptide antagonist of vasopressin V1A and V2 receptors and is clinically used to treat euvolemic and hypervolemic hyponatremia by promoting electrolyte-free water excretion ^33^. Its clinical safety profile makes it an attractive candidate for drug repurposing as a potential antiviral agent. Among the binding poses of Conivaptan in three clusters, the orientation in cluster 1 has the most favorable binding energy (-11.23 kcal/mol). Importantly, Conivaptan also exhibits the strongest predicted binding affinity among all docked molecules. The residue-level interactions involved hydrogen bonds with residues S285, hydrophobic interaction with F204, L239, M262, F242 and stacking interaction with F242 (Figure 3D). Henceforth, Conivaptan was chosen as the most promising small molecule against NiVM and its complex with the representative conformation of cluster1 was selected for subsequent molecular dynamics simulations to assess interaction with residues of the PIP2-binding pocket and membrane-embedded behavior under physiologically relevant conditions.

### 3.3. Dynamics of Conivaptan in the PIP2 binding pocket

To ensure that the docking pose of Conivaptan reflected a genuine binding interaction rather than a static prediction, we performed a 500ns molecular dynamics simulation of the NiVM-Conivaptan complex in explicit solvent. Throughout the simulation, Conivaptan remained stable within the binding pocket, without dissociation or drift into the solvent. The interaction of Conivaptan with the binding pocket residues was monitored over the 500 ns simulation to identify the key interactions responsible for NiVM-Conivaptan binding. The distribution of number of hydrogen bonds between Conivaptan and the polar residues of the binding pocket are shown in Figure 4A(i). One persistent hydrogen bond is observed between Conivaptan and the binding-pocket amino acid GLY240. Additionally, LEU239 also forms backbone-mediated hydrogen bonds with Conivaptan (Figure 4A(ii)). There is also a significant number of hydrophobic interactions between hydrophobic amino acids at the binding pocket and the small molecule (Figure 4B). A comparison of the binding poses of PIP2 and Conivaptan shows that the hydrophobic interacting residues are the same, while the polar interactions differ (Figure 4C). The geometry of the binding interface was preserved, and the ligand-maintained hydrogen-bonding interactions with residues that define the lipid-recognition region. Importantly, the pocket architecture did not collapse or revert toward the apo-like conformation. No unexpected structural perturbations were observed. These results indicate that Conivaptan forms a stable interaction within the PIP2-binding site under dynamic conditions, supporting its progression to membrane-embedded simulations to examine how pocket occupation influences matrix protein-membrane interactions.

**Figure 4:**
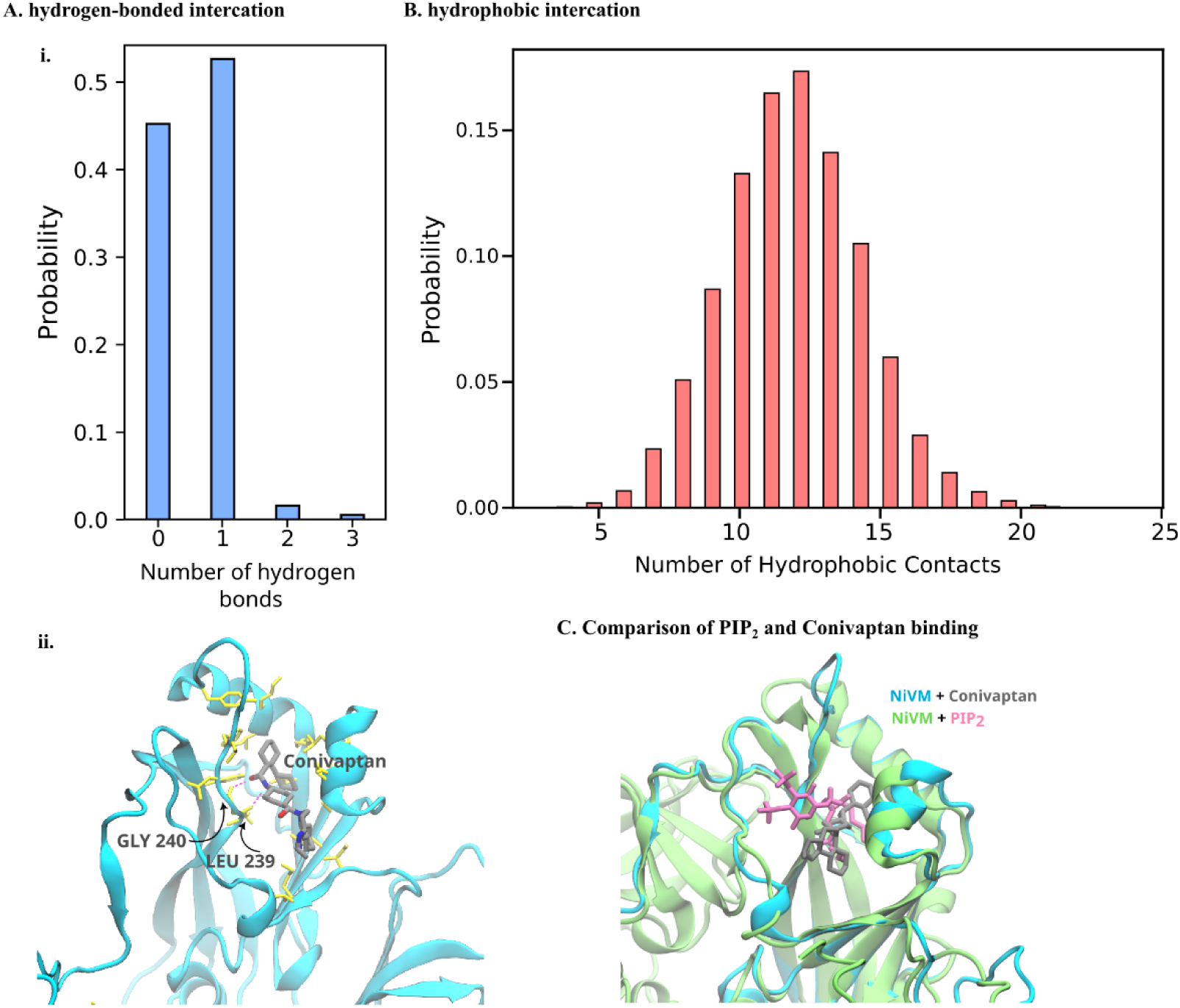
Comparison of the interaction profiles and binding modes of PIP2 and Conivaptan within the NiVM PIP2-binding pocket. (A) (i) Probability distribution of the number of hydrogen bonds formed between Conivaptan and the binding pocket residues during molecular dynamics simulations. (ii) Representative structure of the NiVM-Conivaptan complex highlighting the binding orientation of Conivaptan and the residues involved in hydrogen bond interactions. Hydrogen bonds are shown as dashed lines. (B) Probability distribution of hydrophobic contacts between Conivaptan and the binding pocket residues throughout the simulation. (C) Structural superposition of the NiVM-PIP2 and NiVM-Conivaptan complexes highlighting the relative orientations of the two ligands within the PIP2-binding pocket. Conivaptan occupies a deeper region of the binding cavity while maintaining the overall architecture of the binding pocket.

### 3.4. Effect of ligand binding on membrane interactions of NiVM

The binding of PIP2 to the Matrix protein is followed by its association with the plasma membrane. This induces crucial structural changes in the membrane to induce membrane curvature that eventually leads to virion formation through oligomerization of NiVM-PIP2 at the membrane surface^13^. PIP2-asisted membrane anchoring of other viral proteins like HIV Gag protein and are also reported^34^. The dynamics of Matrix protein at the membrane surface was compared in the absence of any small molecules and in the presence of PIP2 and Conivaptan. The membrane-interacting side of the NiVM dimer is rich in charged residues and are stabilized at the polar head group of the lipid bilayer. The protein-lipid interaction was quantified by calculating occupancies of hydrogen bonds formed between lipid head group and amino acids at the bilayer surface. The number of hydrogen bonds formed between NiVM and bilayer head group increases when there is a ligand bound to the matrix protein binding pocket (Figure 5A). The C-terminal amino acids, including E334, R348 and R351 of NiVM, forming extended interactions with the adjacent monomer in the dimeric state, form multiple hydrogen bonds (Figure 5A(ii-iii)) with the bilayer, which were absent when apo NiVM interacts with the bilayer (Figure 5A(i)). Additional protein-lipid interactions were also found between the N-terminal amino acids (R57, K58, K98 and K142) and bilayer head groups in the presence of ligands. The C-terminal and N-terminal residues are distributed across the protein significantly below the headgroup of bilayer (Figure 5B) in Bilayer_NiVM.

**Figure 5:**
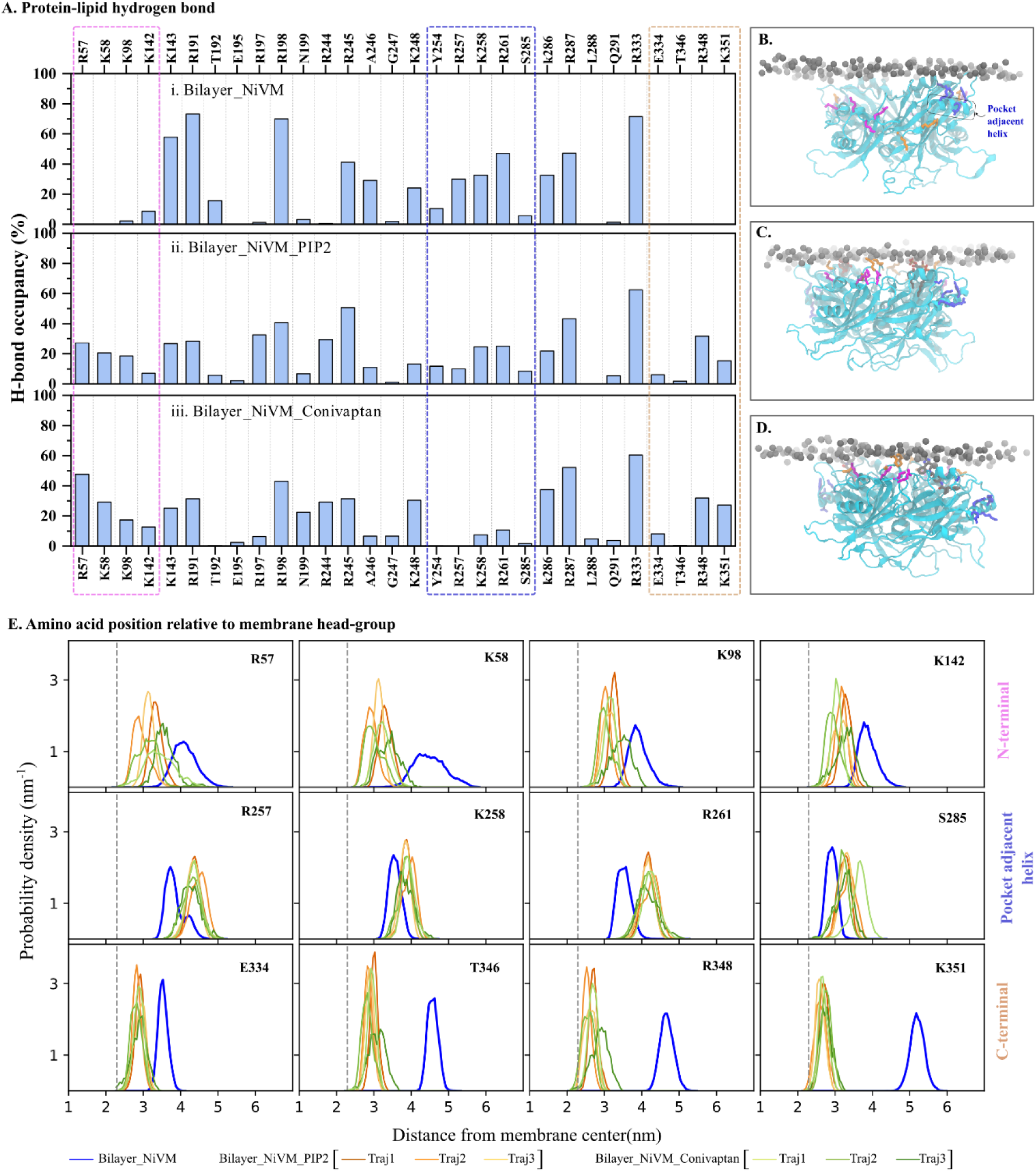
Comparison of protein-membrane interactions of NiVM in the absence and presence of PIP2 or Conivaptan. (A) Hydrogen-bond occupancy between NiVM residues and membrane lipids during simulations of (i) Bilayer_NiVM, (ii) Bilayer_NiVM_PIP2, and (iii) Bilayer_NiVM_Conivaptan. Residues at the N-terminal region, pocket-adjacent helix and C-terminal region are highlighted using magenta, blue and orange rectangles, respectively. Representative snapshots showing the orientation of NiVM at the membrane surface in the (B) Bilayer_NiVM, (C) Bilayer_NiVM_PIP2, and (D) Bilayer_NiVM_Conivaptan systems. Protein is shown as a cyan cartoon, membrane head groups are represented as grey spheres, and residues forming hydrogen bonds with lipid molecules are shown as sticks. (E) Probability distributions of the distances between membrane-interacting residues and the membrane center along the membrane normal for the three systems. Vertical dashed lines indicate the average position of the membrane head-group region and the distribution for Bilayer_NiVM is shown in blue. Independent trajectories of Bilayer_NiVM_PIP2 are shown in red, orange and light orange, while those of Bilayer_NiVM_Conivaptan are shown in yellow, light green and green.

In the presence of PIP2 or Conivaptan, they are clustered below the bilayer headgroup (Figure 5C-D). This relative orientation of these amino acids with respect to headgroups along the membrane normal is shown in Figure 5E. The distances (Z) of bilayer head group phosphate atoms and the amino acid side chains from the bilayer center were calculated, and their distributions are compared. The average Z distance of the bilayer head group from the bilayer center is indicated by the vertical dashed line. The distributions of N-terminal amino acids center approximately around 4 nm in Apo NiVM-bilayer complex. When the ligand-bound NiVM interacts with the bilayer, these distributions shift closer to the bilayer head group, with a mean value of ∼3 nm. Similar reorientations are also observed for C-terminal amino acids E334, T346, R348 and K351 where the mean position of side chain center of mass was shifted by 2-3 nm and overlaps headgroup position. Interestingly, the residues of the helix close to the binding pocket exhibits contrasting behaviour. The occupancies of protein-lipid hydrogen bonds formed by amino acids Y254, R257, R261 and S285 decrease upon ligand binding. Their side chains are distributed closer to the bilayer headgroup in case of Apo NiVM compared to the ligand-bound NiVM (Figure 5E). The difference between bilayer-NiVM interaction in Bilayer_NiVM_PIP2 and in Bilayer_NiVM_Conivaptan is reflected in the occupancies of H-bonded interaction as well as in the type of interacting lipid molecules (Figure S4). There are significant losses in H-bonded interactions in the pocket region residues (Y254, R257, R261 and S285) in the presence of Conivaptan (Figure S4C) compared to PIP2 (Figure S4B).

The loss of interactions is prominent in both monomers in case of Conivaptan in all three trajectories, whereas some interactions are preserved in one monomer while PIP2 is present in the binding pocket. The majority of protein-membrane interactions are mediated through POPI lipids followed by POPC, POPS and PSM. While the interactions in PIP2 are dominated by POPI, there are a significant number of interactions involving POPS and PSM respectively in Conivaptan-bound NiVM.

### 3.5. Effect of ligand binding on membrane physical properties

The alteration in protein-membrane interaction when different small molecules bind to the NiVM pocket had affected the physical properties of lipid-bilayer. Membrane thickness is measured as the average distance between the phosphate groups of the upper and lower leaflet. Thickness is calculated at different bins across the bilayer plane (Figure 6A). A lower inter-head-group distance indicates membrane thinning, while a larger distance indicates a relatively thicker membrane. The Bilayer in absence of any binding partner is the thickest among all systems and the thickness vary from 4.8nm to 5nm (Figure 6A(a)). Upon attachment of NiVM on the surface, there is a significant drop of ∼0.3 nm in the thickness map (Figure 6A(b)). The decrease in membrane thickness is due to interactions with the surface residues of NiVM. Interestingly, the uniformity in the Thickness map is maintained in Bilayer_NiVM, similar to Bilayer, despite the overall thinning of the membrane. In the presence of PIP2 or Conivaptan, there are significant changes in the thickness value and its distribution across the membrane surface. To understand the alteration in thickness relative to the ligand-binding pocket, the positions of the centers of mass of PIP2 or Conivaptan are mapped on the thickness map as red dots. In the presence of PIP2, there is a significant drop in membrane thickness, specifically around the PIP2-binding pocket (Figure 6A(c)). The extent of membrane thinning is less while Conivaptan is bound to the PIP2-binding pocket. The thickness is significantly higher around the binding pocket in Bilayer_NiVM_Conivaptan (Figure 6A(d)). There is significant membrane thickening around the binding pocket and the overall thickness is even higher than Bilayr_NiVM. The observations are consistent in the three replicates of both Bilayer_NiVM_PIP2 (Figure 6A(c(i-iii))) and Bilayer_NiVM_Coniaptan (Figure 6A(d(i-iii))). The absence of membrane thinning around the binding pocket in the presence of Conivaptan compared to that in the presence of PIP2 can be attributed to the reduced interaction of helix adjacent to binding pocket. The significant alteration of local membrane thickness can affect the overall compactness of the arrangement of lipid molecules in the bilayer. This compactness is measured by the average area per lipid molecule (APL). The distribution of APL, calculated over the simulation, is compared for the systems and their replicates (Figure 6B).

**Figure 6:**
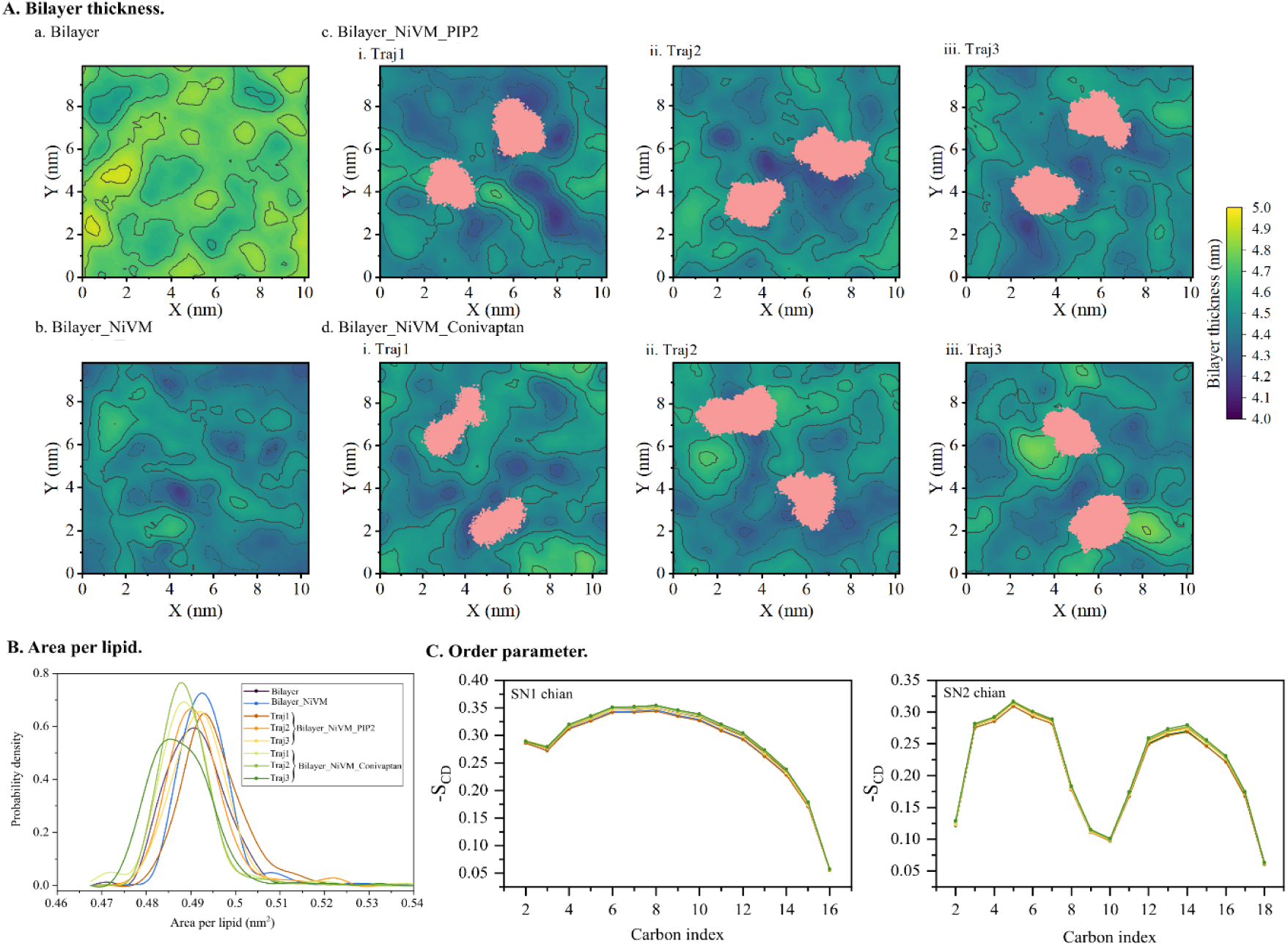
Comparison of membrane deformation and lipid packing induced by NiVM in the absence and presence of PIP2 or Conivaptan. (A) Bilayer thickness maps for (a) the lipid bilayer alone, (b) Bilayer_NiVM, (c) Bilayer_NiVM_PIP2 (three independent trajectories), and (d) Bilayer_NiVM_Conivaptan (three independent trajectories). Positions of PIP2 or Conivaptan projected on the membrane surface (X-Y plane) during simulation are shown by pink dots. (B) Probability density distributions of the area per lipid for the bilayer alone and membrane systems containing NiVM, NiVM– PIP2, and NiVM–Conivaptan. (C) Deuterium order parameters (−SCD) of the phospholipid SN1 and SN2 acyl chains for the different simulation systems. Independent trajectories of Bilayer_NiVM_PIP2 are shown in red, orange and light orange, while those of Bilayer_NiVM_Conivaptan are shown in yellow, light green and green.

The APL for PIP2-bound NiVM and its replicas are higher compared to the APL for Conivaptan-bound NiVM. Similar to membrane thickness, the APL of Bilayer_NiVM_Conivaptan is significantly different from that in Bilayer_NiVM. This clearly demonstrates the difference in membrane compactness when the natural substrate PIP2 is replaced by the inhibitor molecule Conivaptan. The arrangement of lipid molecules in the bilayer is also quantified using acyl chain order parameter (S_CD_). Order parameters were calculated separately for SN1(saturated Palmitoyl chain) and SN2 (unsaturated Oleyl chain) chains (Figure 6C). For both SN1 and SN2 chains, the order parameter value increases when Conivaptan is bound to NiVM compared to the PIP2-bound NiVM. The higher orderliness of bilayer is consistent in the three replicate trajectories. The increase in order parameter in the presence of Conivaptan can be attributed to the lesser membrane deformation due to reduced protein-membrane interaction.

## 4. Discussion

The essential role of the Nipah virus matrix protein in viral assembly and membrane association has made it an attractive target for antiviral drug discovery. Consequently, several computational studies have explored small-molecule inhibitors targeting different functional regions of the protein. For example, Macalad B. et. al. targeted matrix protein using different metabolites of *Streptomyces* spp. and identified several small molecules as potential inhibitor binding at the substrate region and dimerization region^35^. Yang et. al. used deep learning coupled with MD simulation to identify Rutin and Lactitol as matrix protein inhibitor^36^. While these studies provide valuable insights into ligand recognition by the NiVM binding pocket, they primarily evaluate inhibitor binding using a single receptor conformation and do not consider the impact of inhibitor binding on the membrane-associated form of the matrix protein. Furthermore, the conformational flexibility of the PIP2-binding pocket, which may influence ligand recognition and inhibitor binding, has largely been overlooked. These limitations are addressed in the present study through the integration of receptor conformational dynamics with membrane-associated functional analyses.

The MD simulation of the PIP2-bound NiVM has revealed the dynamic binding of PIP2 at the NiVM pocket. The binding modes differ in terms of the conformation of the binding pocket and the corresponding protein-ligand interactions. In majority of the conformations PIP2 exhibit few or no hydrogen bonds with the pocket residues whereas a smaller fraction of conformations forms with one or more hydrogen bonds (Figure 2A(i)). The hydrophobic contacts range from 10 to 20 with an average of ∼16 hydrophobic contacts (Figure 2B(i)). Together the variability in hydrogen bonding and hydrophobic contacts underscore the conformational plasticity of the PIP2-binding pocket. To account for this conformational heterogeneity, the three most populated clusters were selected for ensemble-based virtual screening, leading to the identification of Conivaptan as the top-ranked compound capable of binding all representative pocket conformations (Table S1). A comparison of binding of PIP2 and Conivaptan to the NiVM pocket reveals more persistent hydrogen-bonded interaction while maintaining a comparable hydrophobic contacts (Figure 3A-B). The smaller molecular size also enables Conivaptan to penetrate deeper into the pocket (Figure 3C) potentially contributing to the enhanced polar interaction with surrounding residues. Collectively, these observations demonstrate that accounting for binding-pocket conformational heterogeneity enables the identification of inhibitors capable of compact binding across multiple receptor conformations.

The importance of PIP2 binding to the NiVM pocket for its interaction with plasma membrane lies on the large conformational change of the C-terminal amino acids (Q328-K351) evident from the crystal structure (Figure 1(B-C))^13^. These conformational changes lead to formation of multiple polar interactions with lipid head groups involving E334, R348 and K351 (Figure 5A) resulting in stronger association at the membrane surface. In addition to this we have identified additional interaction of the N-terminal residues R57, K58, K98 and K142 with lipid molecules when PIP2 is bound to NiVM (Figure 5A). Although the initial orientations of these residues are similar in crystal structure, the conformational changes of surrounding residues favor stronger interaction during the course of simulation. Overall, the introduction of PIP2 in the NiVM pocket has rearranged the surface electrostatic towards favorable electrostatic interaction with bilayer. The membrane-protein interaction is altered when PIP2 in the binding pocket is replaced by Conivaptan. There is a significant drop in the interaction of the residues of pocket adjacent helix e.g. Y254, R257, K258, R261 and S285 (Figure 5A). Conivaptan has also affected the types of lipid molecules NiVM interacts with. The major interactions with POPI are replaced by interactions with POPS and PSM (Figure S4). These changes indicate that PIP2 binding strengthens the membrane-binding interface of NiVM, whereas its replacement by Conivaptan reorganizes the protein–membrane interactions and alters the mode of membrane association.

The altered protein-membrane interactions are accompanied by pronounced changes in the physical properties of the surrounding lipid bilayer. Influenza A virus matrix protein has been shown to induce membrane curvature and deformation on negatively charged membranes, highlighting the ability of viral matrix proteins to remodel host membranes.^37^ Consistent with this general behavior, our simulations show that NiVM induces membrane thinning around the protein-contacting regions. NiVM binding alone induces moderate membrane thinning, whereas PIP₂ binding enhances this effect by promoting a more localized deformation centered around the PIP₂-binding pocket. Replacement of PIP2 by Conivaptan substantially alters the membrane thickness profile. Compared with the PIP2-bound system, the membrane exhibits a higher average bilayer thickness, with some replicas showing localized thickening beneath the protein region surrounding the binding pocket. This difference is further reflected in the area per lipid and acyl-chain order parameters. The lower area per lipid together with the higher acyl-chain order indicates a more compact and ordered membrane^38^, consistent with the reduced membrane deformation observed in the presence of Conivaptan compared with PIP2.

The conformational analysis of NiVM in aqueous solution and at the plasma membrane reveal two complementary mechanisms underlying the inhibitory effect of Conivaptan. In solution, Conivaptan competitively replaces PIP2 and forms stronger polar interaction with the binding pocket residues. At the membrane, however, Conivaptan reorganizes the protein-membrane interactions, resulting in reduced membrane deformation. This suppresses membrane thinning and lipid reorganization, that are likely required for the subsequent membrane bending events associated with virion assembly. Together, these findings suggest that Conivaptan stabilizes a membrane-binding state of NiVM that is less competent for membrane remodeling, thereby impairing the early membrane-associated steps involved in viral assembly.

## Conclusion

In summary, the present study provides a molecular perspective on how competitive inhibition of the Nipah virus matrix protein extends beyond ligand displacement at the PIP2-binding pocket. By incorporating the conformational heterogeneity of the binding site through ensemble docking and analyzing protein dynamics in both aqueous and membrane environments, we demonstrate that the FDA-approved drug Conivaptan binds more favorably than the natural ligand PIP2 while altering the membrane-associated functions of the matrix protein. Unlike previous approaches that primarily focused on binding affinity, our analyses reveal that inhibitor binding reorganizes protein-membrane interactions, resulting in reduced membrane thinning, altered lipid organization, and diminished membrane-remodeling capability. Together, these findings support a dual mechanism of inhibition in which Conivaptan not only competes with PIP2 for binding but also attenuates the membrane-remodeling activity required for efficient viral assembly and budding. More broadly, this study highlights the importance of considering receptor conformational dynamics together with membrane-associated functional consequences during antiviral drug discovery and provides a framework for the rational design and evaluation of next-generation inhibitors targeting the Nipah virus matrix protein. The mechanistic insights presented here provide a strong foundation for future biochemical studies to evaluate Conivaptan and related inhibitors targeting the Nipah virus matrix protein.

## Data availability

The coordinates of top ten ligands for the three clusters and the starting structure and parameter files used for MD simulation have been deposited in Zenodo repository and are publicly available at https://doi.org/10.5281/zenodo.21383342.

## Acknowledgement

APK and AM are thankful to the Department of Biotechnology (DBT), Govt. of India for their fellowships. Authors are thankful for funding from the Department of Biotechnology (DBT), Govt. of India (grant no. BT/PR40175/BTIS/137/41/2022) for the Bioinformatics Centre. Authors acknowledge the Param Shakti supercomputing facility of IIT Kharagpur established under National Supercomputing Mission (NSM), Government of India supported by Centre for Development of Advanced Computing (CDAC), Pune. APK acknowledges Prof. Saraboji Kadhirvel of Central University of Punjab for valuable discussions.

## Author contributions

APK performed the simulations, formal analysis and wrote the initial draft of the manuscript. AM conceptualized the work, designed the simulation and performed the formal analyses. RPB supervised the work and arranged funding and resources. All authors approved the final version of the manuscript.

