## Supporting Information for "Competitive Inhibition of the Nipah Virus Matrix Protein by Conivaptan: From Binding Pocket Dynamics to Protein-Membrane Interactions"

for

**Figure S1**

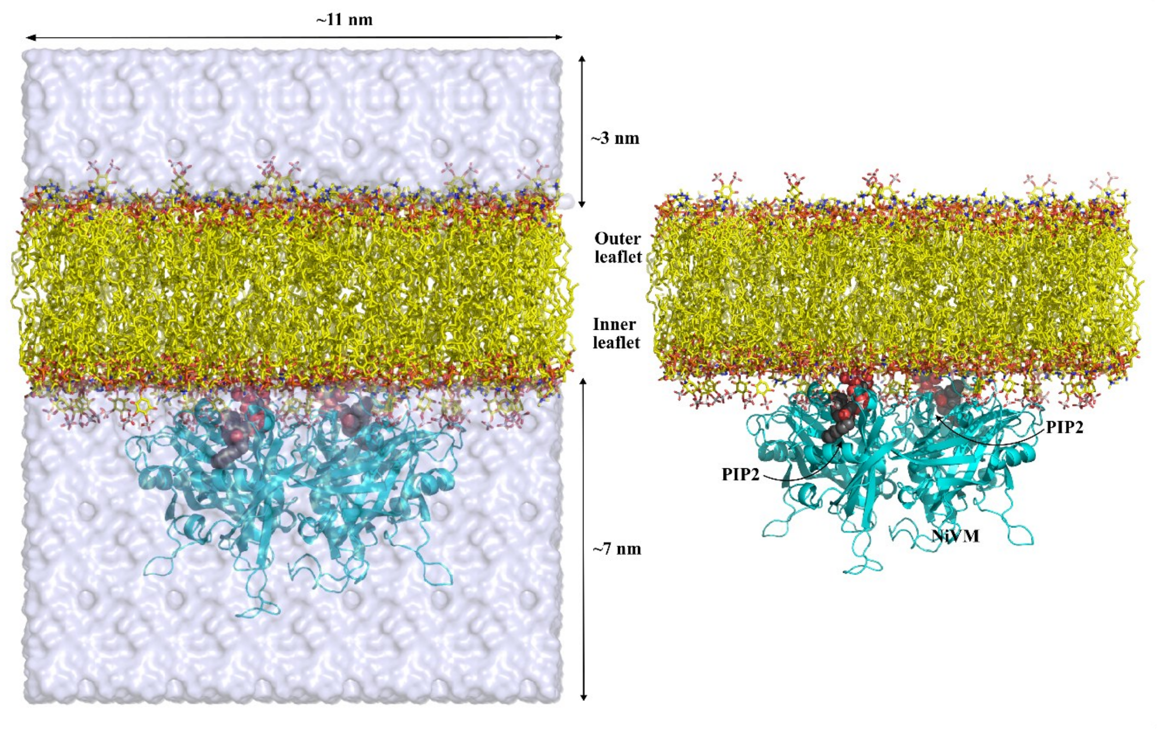


**Figure S1:** Different components of protein-membrane assembly. Lipid molecules are shown in yellow stick. The oxygen and phosphorus atoms are coloured red and orange, respectively. The water layer on both side of the membrane are shown as light blue surface. The Matrix protein is shown as cyan cartoon while the PIP2 molecules at the protein-membrane interface are shown in grey spheres. The system dimensions are also labelled.

**Figure S2**

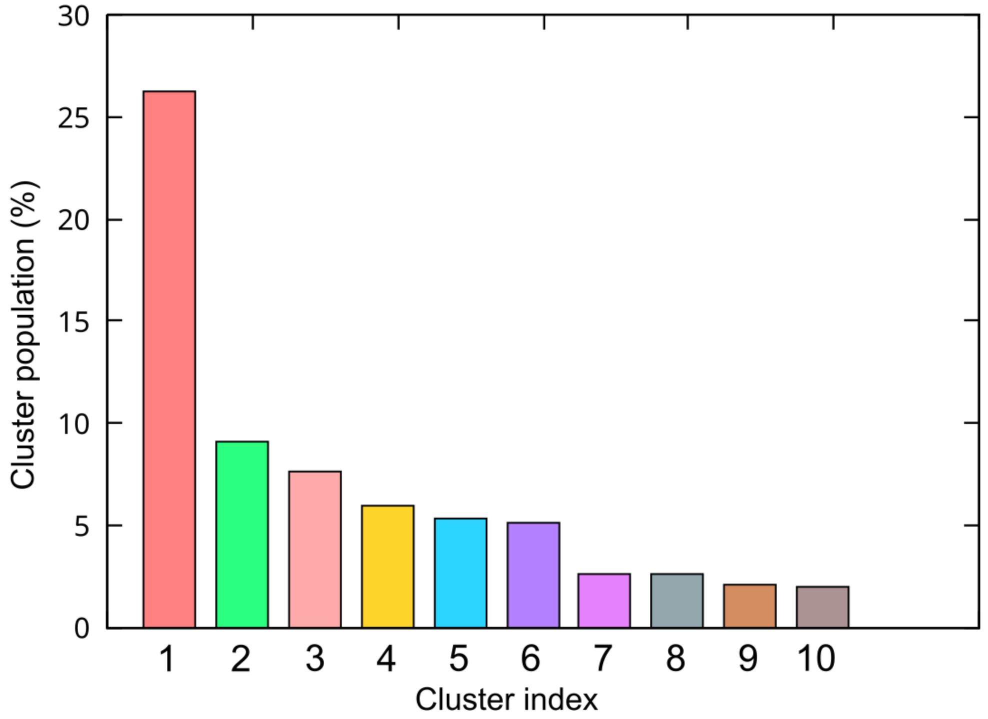


**Figure S2:** Population of major clusters obtained from the clustering of the trajectory of NiVM_PIP2 based on the RMSD of binding pocket amino acids.

**Figure S3**

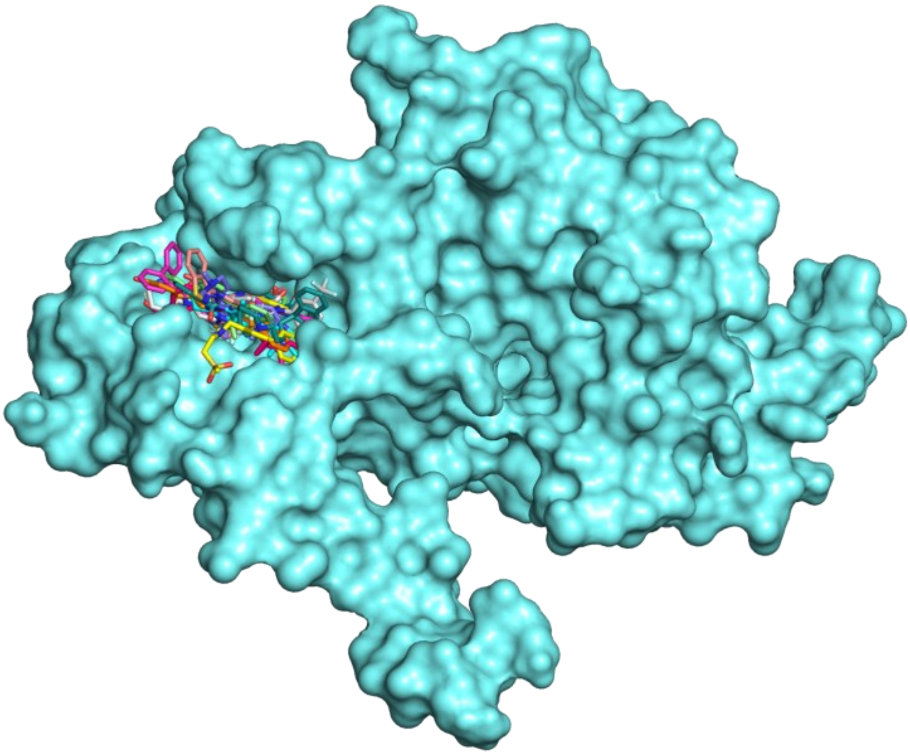

**Figure S3:** Overlay of the top ten small molecules targeting NiVM pocket of cluster1 conformation.

### Figure S4

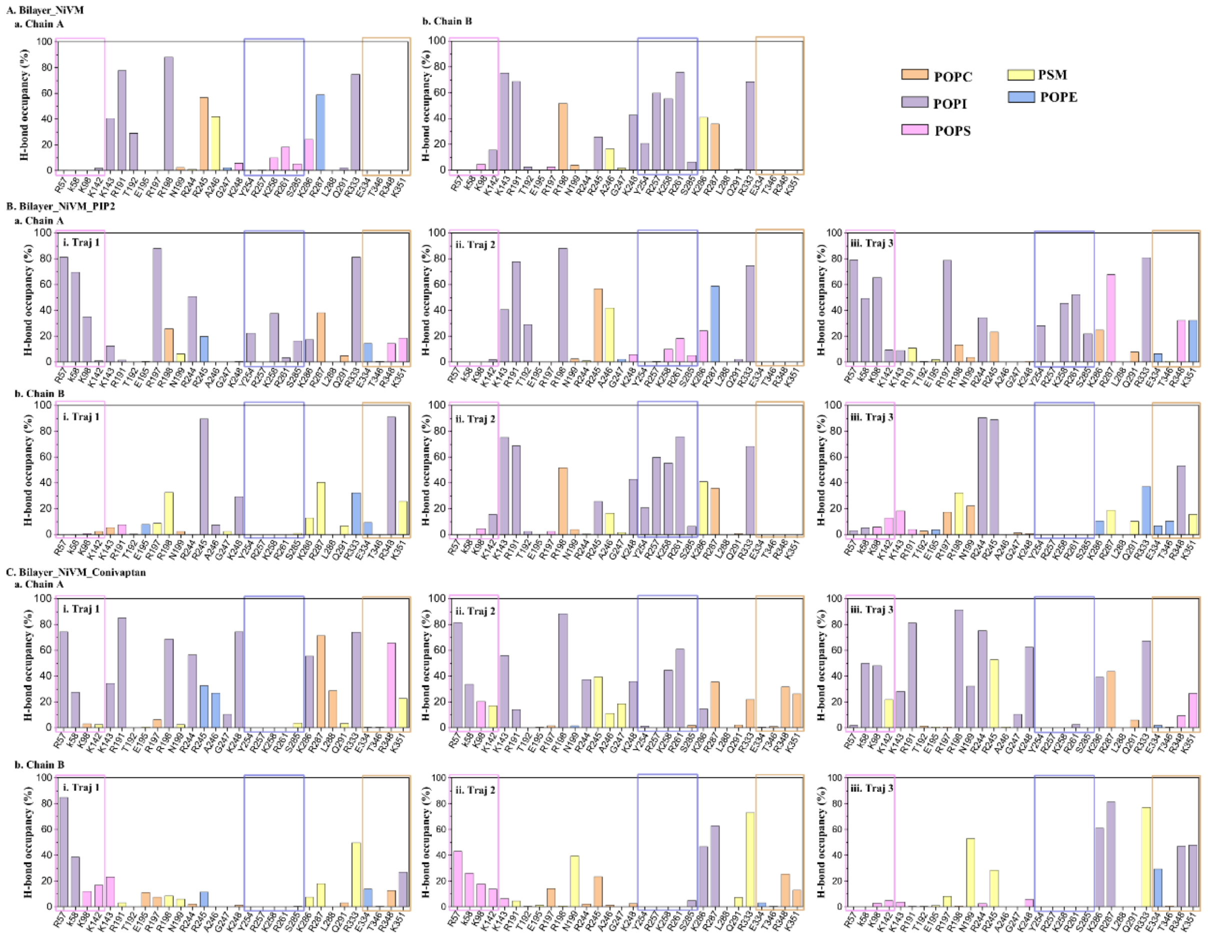


**Figure S4:** Protein-lipid hydrogen-bond interactions of the Nipah virus matrix protein (NiVM) in different membrane-bound systems. (A) Hydrogen-bond occupancy between NiVM and membrane lipids in the Bilayer_NiVM system for (a) Chain A and (b) Chain B. (B) Hydrogen-bond occupancy for the Bilayer_NiVM_PIP2 system showing three independent trajectories (i–iii) for (a) Chain A and (b) Chain B. (C) Hydrogen-bond occupancy for the Bilayer_NiVM_Conivaptan system showing three independent trajectories (i–iii) for (a) Chain A and (b) Chain B. Bar colors represent different lipid species (POPC, POPI, POPS, PSM, and POPE). Colored boxes highlight residues belonging to the N-terminal region (magenta), the binding pocket adjacent helix (blue), and the C-terminal region (orange). Hydrogen-bond occupancy was calculated as the percentage of simulation frames in which a residue formed at least one hydrogen bond with the corresponding lipid species.

**Table S1:** List of top 10 small molecules targeting three clusters obtained from Autodock vina. The predicted affinity values (in kcal/mol), chemical name and Pubchem CID are provided for each molecule.

| Index | Cluster 1 | | | Cluster 2 | | | Cluster 3 | | |
|---|---|---|---|---|---|---|---|---|---|
| | Name | Pubchem CID | Affinity | Name | Pubchem CID | Affinity | Name | Pubchem CID | Affinity |
| 1 | Conivaptan | 151171 | -11.23 | Conivaptan | 151171 | -10.89 | Conivaptan | 151171 | -10.44 |
| 2 | Telmisartan | 65999 | -9.88 | Midostaurin | 9829523 | -10.76 | Adapalene | 60164 | -10.06 |
| 3 | Midostaurin | 9829523 | -9.81 | Dihydroergotamine | 10531 | -10.58 | pimozide | 16362 | -9.63 |
| 4 | Dutasteride | 6918296 | -9.78 | Ergotamine | 8223 | -10.32 | Dutasteride | 6918296 | -9.47 |
| 5 | Lurasidone | 213046 | -9.73 | Irinotecan | 60838 | -9.81 | Palbociclib | 5330286 | -9.26 |
| 6 | Lapatinib | 208908 | -9.60 | Lomitapide | 9853053 | -9.75 | Rolapitant | 10311306 | -8.94 |
| 7 | Lomitapide | 9853053 | -9.6 | Adapalene | 60164 | -9.74 | Dihydroergotamine | 10531 | -8.86 |
| 8 | Nilotinib | 644241 | -9.53 | Tipranavir | 54682461 | -9.64 | Aprepitant | 135413536 | -8.78 |
| 9 | Lumacaftor | 16678941 | -9.55 | Bromocriptine | 31101 | -9.59 | Lumacaftor | 16678941 | -8.76 |
| 10 | Netupitant | 6451149 | -9.49 | Nilotinib | 644241 | -9.45 | Midostaurin | 9829523 | -8.73 |
